# The population and evolutionary dynamics of bacteriophage: Why be temperate revisited

**DOI:** 10.1101/824235

**Authors:** Waqas Chaudhry, Nicole Vega, Adithi Govindan, Rodrigo Garcia, Esther Lee, Ingrid McCall, Bruce Levin

## Abstract

Bacteriophages are deemed either lytic (virulent) or temperate, respectively depending on whether their genomes are transmitted solely horizontally, or both horizontally and vertically. To elucidate the ecological and evolutionary conditions under which natural selection will favor the evolution and maintenance of lytic or temperate modes of phage replication and transmission, we use a comprehensive mathematical model of the dynamics of temperate and virulent phage in populations of bacteria sensitive and resistant to these viruses. For our numerical analysis of the properties of this model, we use parameters estimated with the temperate bacteriophage Lambda, λ, it’s clear and virulent mutants, and *E. coli* sensitive and resistant - refractory to these phages. Using batch and serial transfer population dynamic and reconstruction experiments, we test the hypotheses generated from this theoretical analysis. Based on the results of this jointly theoretical and experimental study, we postulate the conditions under which natural selection will favor the evolution and maintenance of lytic and temperate modes of phage replication and transmission. A compelling and novel prediction this *in silico*, *in vitro*, and *in plastico* study makes is lysogenic bacteria from natural populations will be resistant-refractory to the phage for which they are lysogenic as well as lytic phage sharing the same receptors as these temperate viruses.

## Introduction

From the perspective of population and evolutionary biology, the single most important functional distinction among bacteriophage (phage) is whether they are <u>lytic</u> (virulent) or temperate. Lytic phages are exclusively transmitted horizontally by infecting susceptible bacteria within which they replicate and from which they are released into the environment, most commonly with the demise of the infected bacterium. In addition to this capacity for horizontal transmission, temperate phage can also incorporate their genomes into that of the infected cell, as a <u>prophage</u> [1] and be transmitted vertically, in the course of cell division by these <u>lysogenic</u> bacteria [2–4].

On first consideration, it would seem that in virtually all environments, natural selection would favor a temperate rather than a purely lytic mode of phage replication and transmission. 1-Upon invading a new population of sensitive bacteria, temperate phage are transmitted primarily horizontally and would sweep through the susceptible population of bacteria at rates similar to that of purely lytic phage with the same infection parameters, adsorption rates, latent periods and burst sizes [5]. 2-By establishing lysogens, the genomes of temperate phages are transmitted vertically and the phage can be maintained in the bacterial population when the densities of their host bacteria are too low for these viruses to persist by horizontal transmission alone [3, 6–9]. 3-The prophage of temperate viruses can carry genes that in some environments augment the fitness of their host bacteria and thereby the fitness of the bacteriophage (specifically virulence factors [10–12] and drug-resistance cassettes [13, 14], but including genes whose functions have not been identified [15, 16]). 4-By induction and release of free phage, lysogeny can provide an advantage to its host population by allelopathy, killing members of susceptible competing populations [17–21].

There are, however, two major caveats to these arguments about a universal advantage of a temperate mode of phage replication and transmission relative to a purely lytic mode, both based on the conditions under which lytic phage might be able to invade a temperate population. (1) through mutation, temperate phage can become lytic, [1, 22, 23, 24, 25] and kill lysogens [26] as well as sensitive non-lysogens. If the densities of the bacteria are great enough to maintain phage by horizontal transfer alone, these virulent mutants may eliminate their temperate ancestors [5]. (2) through mutation, bacteria can become resistant and refractory to infections with both temperate and lytic forms of a given phage. If this envelope resistance arises in lysogens, free phage will be generated by spontaneous induction, leading to maintenance of the temperate phage population even in the absence of new infections. If resistance arises in non-lysogens in the same environment, these cells would be invulnerable to infection by lytic and temperate forms of the phage, and resistant cells would be anticipated ascend to dominate the bacterial population. Unless there is some other mechanism for maintaining the phage, like physical refuges [27] or leaky resistance [28], the phage would be lost. Thus, the answer to the “why be temperate” question ultimately comes down to elucidating (i) the conditions under which a temperate mode of phage replication can be maintained in situations where virulent mutants are present or can be generated, and (ii) the conditions under which the resistant cells that evolve will be lysogens or not.

In this study, we use a simple, but comprehensive mathematical model and computer simulations to elucidate the *a priori* conditions under which temperate phage, their clear and lytic mutants will become established and maintained in populations of sensitive bacteria in the absence and presence of resistant mutants. For the numerical analysis of properties of this model in simulations, we use estimates of the parameters obtained with *E. coli* K12 and the phage Lambda (λ). To explore the validity and limitations of this model and test the predictions made from our analysis of its properties, we perform population dynamic experiments with the phage λ, its clear and lytic mutants, and *E. coli* that are sensitive and resistant to these viruses. Based on the results of this jointly theoretical and experimental analysis, we postulate the conditions under which selection will favor the evolution and maintenance of temperate modes of phage replication.

## RESULTS

### I Theoretical (*a priori*) Considerations

#### A comprehensive mass-action model of the population dynamics of temperate phage and lysogeny

The model developed here is an extension of that in [8] and [28] (Fig 1). There are five populations (states) of bacteria: sensitive non-lysogens (N), lysogens (L), newly formed lysogens (N_L_), resistant lysogens (N_LR_), and resistant non-lysogens (N_R_) (cells per ml); and three populations of free phage: temperate (P) lytic (P_v_), and clear (P_c_) (particles per ml). The clear mutant P_C_ can replicate on sensitive cells but can neither form lysogens nor replicate on lysogens. Resistant non-lysogens are generated by mutation from sensitive cells, (S--> N_R_) and resistant lysogens are generated from new lysogens (N_L_ -- >N_RL_). Temperate phage generate clear mutants (P --> P_C_) and virulent mutants (P_C_ --> P_V_) [25].

**Fig 1.**
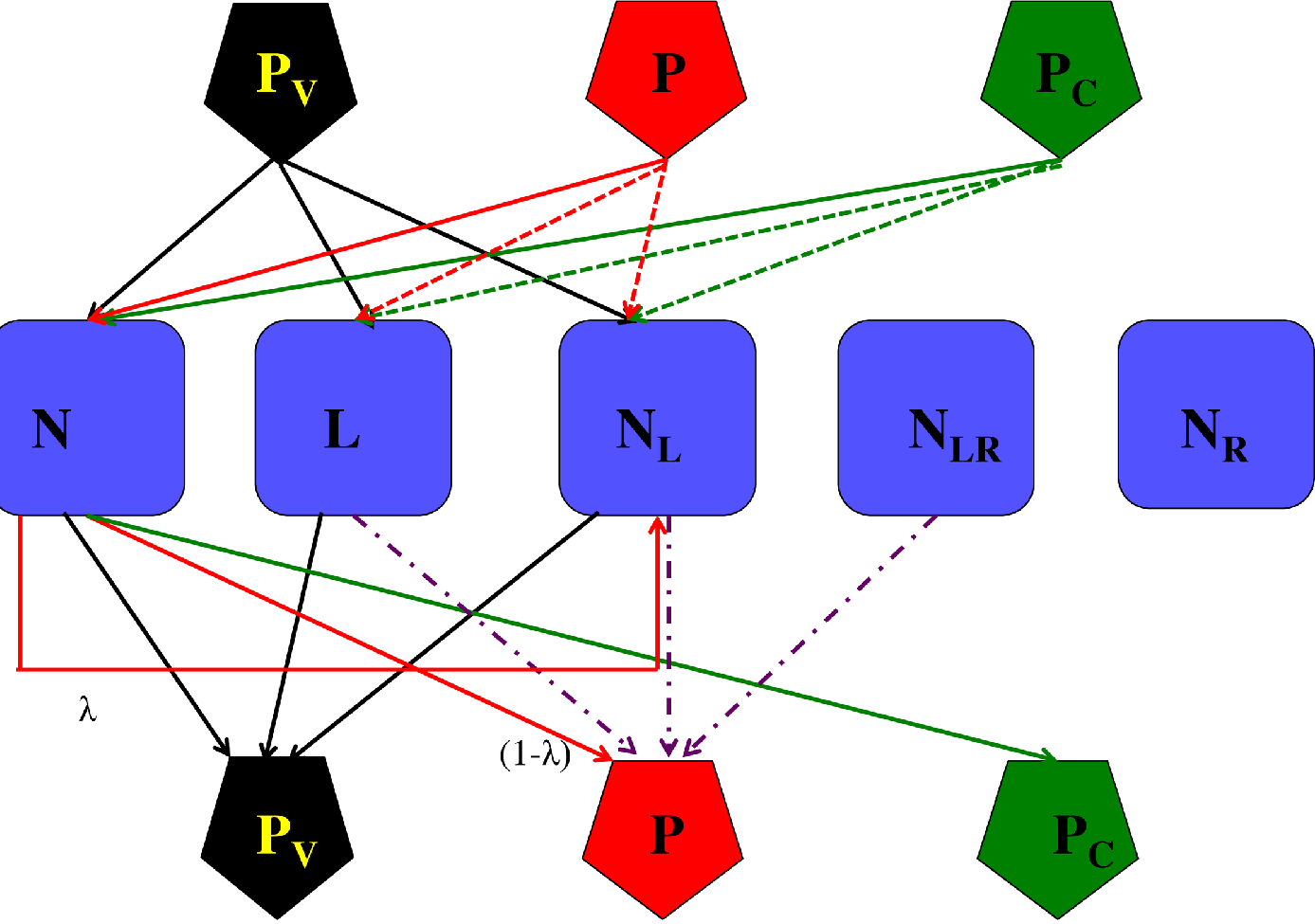
A comprehensive model of the population and evolutionary dynamics of temperate, P, clear P_C_ and virulent, P_V_ phage with sensitive, N, lysogen, L, newly formed lysogens, N_L_, resistant lysogens, N_LR_ and resistant non-lysogens, N_R_. Solid thin lines - adsorption and lytic production of phage of the noted type. Broken lines-adsorption and loss of the phage due to immunity. Dot - dash lines-production of phage by induction of lysogens. With lytic infection and induction, the cell is killed. A fraction λ of the infections of the of sensitive cells N, by temperate phage, P, produce new lysogens, N_L_, the remaining (1-λ) of these infections are lytic.

These bacteria grow at maximum rates *vn*, *vl*, *vnl*, *vlr* and *vr* per cell per hour respectively, with their net rate of growth being equal to the product of their maximum growth rates and Ψ(r)=R/(R+k), where R, μg/ml is the concentration of the limiting resource and k (μg/ml) is the concentration of the limiting resource at which the net growth rate is half its maximum value, the Monod constant [29]. The phage adsorb to the bacteria at a rate proportional to the product of their densities, an adsorption rate parameter, δ, (per hour per ml) and the resource function Ψ(r). The latter is based on the assumption that as the resources decline, the rates of replication of the phage decline, and when there are no resources, the phage can no longer infect the bacteria. A fraction of the adsorptions λ(0 < λ < 1) of free temperate phage *P* to sensitive non-lysogens *N* generate newly formed lysogens *N_L_*, with the remaining (1-λ) infections being lytic. In this model, we are assuming that the probability of lysogen formation is independent of the relative densities of phage and sensitive hosts, the multiplicity of infection, MOI.

All the phage can infect the sensitive cells, but only P can form lysogens, which are immune to infections by P and clear mutants, P_C_. When free temperate phage or clear mutants, P, and P_C_ adsorb to the lysogens, *L* and *N_L_*, the phage are lost and removed from their respective populations. When the lytic phage, P_V_ adsorb to sensitive cells or lysogens (N, L or N_L_) they kill those bacteria and replicate. The resistant bacteria, NR and NLR, are refractory to all the phage; they are not adsorbed.

At a rate *i* per cell per hour, lysogens, L, N_L_ and N_LR_ are induced, die and produce β (burst size) free phage particles. This same number of phage, β, is also produced by 1-l (0 < l < 1) of adsorptions of P to N that result in lytic infection as well as infections with virulent phage, P_V_ on sensitive cells and lysogens, L and N_L_. Clear phage, P_C_, replicate on sensitive non lysogens, N, producing b free phage, but are lost via sorption when they infect lysogens, N and N_L_. We neglect the latent period and the loss of phage due to intrinsic mortality. As with our lytic phage model [28], we assume there are transition rates at which resistant bacteria revert to their corresponding sensitive states, N_R_→N, N_LR_→N_L_ respectively *μrs* and *μrl*. There are also transition rates, *μsr* and *μlr*, at which sensitive bacteria generate their corresponding resistant states, N→N_R_ and NL→N_LR_. Finally, we assume there is a mutation rate, μ, at which temperate phage generates clear or virulent mutants, P → P_C_ and P → P_V_. All of these rates decline with the concentration of the limiting resource R, via the Ψ(R) function. The variables and parameters of this model are separately defined and the range of values used for the parameters are presented in Table 1.

**Table 1 –.** Variables and Parameters of a model of the population and evolutionary dynamics of bacteriophage.

| Variable | Definition* |
| --- | --- |
| N | Sensitive, non-lysogens - can form lysogens |
| L | Lysogens (initially present) |
| N<sub>L</sub> | New lysogens (generated in the course of the experiment) |
| N<sub>R</sub> | Resistant non-lysogens (refractory to all phage) |
| N<sub>LR</sub> | Resistant lysogens (refractory to all phage, but carry the prophage) |
| P | Free temperate phage (can infect N and form lysogens) |
| $P_c$ | Free clear phage (can infect N, but not form lysogens) |
| $P_v$ | Free virulent phage (can infect all but resistant cells, $N_R$ and $N_{LR}$.) |
| $r$ | The concentration of the limiting resource ($\mu\text{g/ml}$). |

\*All densities in cells or particles per ml

| Parameter | Definition and dimensions | Range of values |
| --- | --- | --- |
| $e$ | Conversion efficiency $\mu\text{g/cell}$ | $5\text{E-}7$ |
| $k$ | Monod constant - $\mu\text{g/ml}$ | 1 |
| $v$ | Maximum growth rate sensitive, N per cell per hour | 0.7- 1.2 |
| $v_L$ | Maximum growth rate lysogens, L, $N_L$ per cell per hour | 0.7-1.2 |
| $v_r$ | Maximum growth rate resistant, $N_R$, $N_{LR}$ per cell per hour | 0.7 -1.2 |
| $\delta$ | Adsorption rate constant ml per cell per phage per hour | $2\text{E-}7$ |
| $\beta$ | Burst size, particles per infected sensitive cell | 50 |
| $\beta_L$ | Burst size, particles per induced lysogen | 50 |
| $\lambda$ | Probability of lysogeny upon infection of N | $10^{-1} \text{ -- } 10^{-5}$ |
| $i$ | Rate of induction - per lysogen per hour | $10^{-3} \text{ -- } 10^{-5}$ |
| $\mu_{sr}$ | Transition rate $S \rightarrow N_R$ and $N_L \rightarrow N_{LR}$ per cell per hour | 0 or $5 \times 10^{-5}$ |
| $\mu_{rs}$ | Transition rate $N_R \rightarrow S$ and $N_{LR} \rightarrow N_R$ per cell per hour | 0 or $5 \times 10^{-45}$ |
| $\mu$ | Rate of mutation $P \rightarrow P_c$ and $P \rightarrow P_v$ per cell per hour | 0 or $10^{-8}$ |

With these definitions and assumptions, the rates of change in the densities of the bacteria and phage and concentration of resource are given below by the set of differential equations.

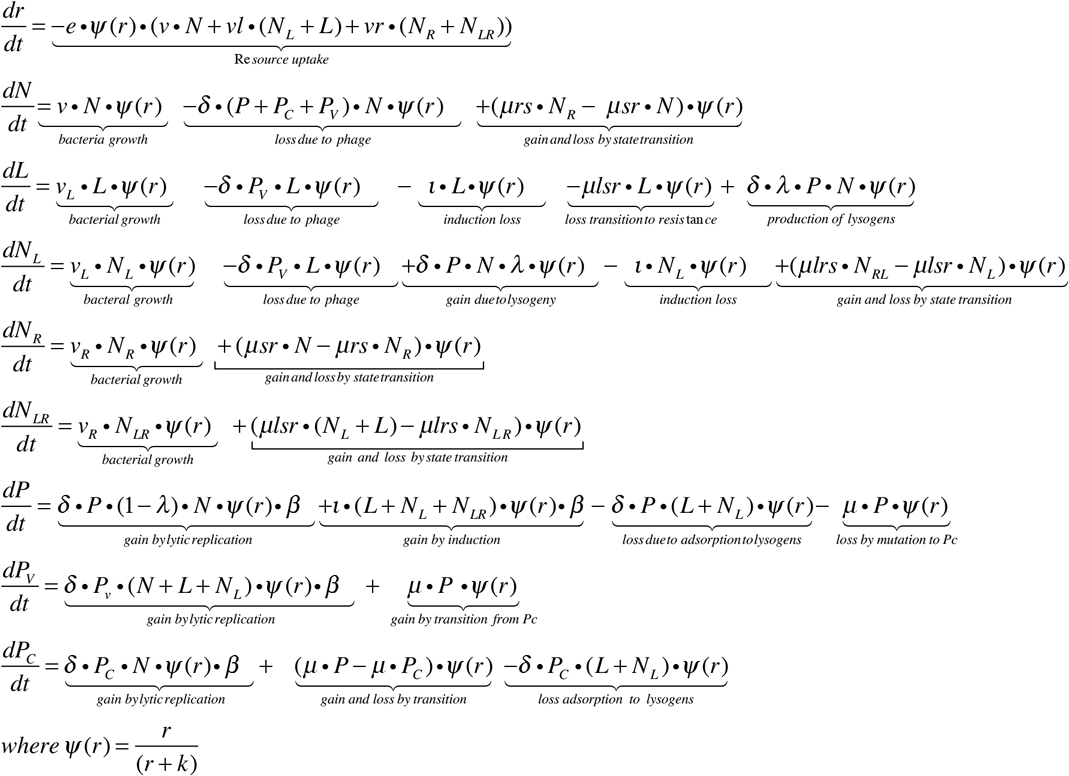

#### Numerical Analysis – computer simulations

For our exploration of the properties of this model we use numerical solutions programmed in Berkeley Madonna^™^. To be consistent with the experiments, for the longer-term population dynamic simulations, we consider the maintenance of the community by serial transfer; every 24 hours, all components are diluted by a factor of *d* (0<d<1) and *C* μg/ml of fresh resource are added. Unless otherwise noted, the values of the parameters used for these simulations are in the range estimated in [28, 30] or in this study (see Materials and Methods).

### Theoretical Results

#### 1-Simulated population dynamics in the absence of resistance

In Fig 2 we follow the changes in the densities of these bacterial and phage populations when resistance does not develop in the bacterial populations (μsr =μlsr = 0). In this and the following simulations, the concentration of the limiting resource at the start of each transfer is R (0) = C= 250 μg/ml. With this concentration of maltose, in the absence of phage the density of the bacteria at the end of each transfer (5 ×10^8^ cells per ml) is similar to that in the maltose limited M9 minimal medium used in our experiments. In the absence of virulent phage, whether temperate phage are introduced as free phage, P, (Fig. 2A) or as lysogens, L, (Fig 2 B), within short order the population is dominated by “new” lysogens (NL). In the resulting populations, lysogens are then maintained at the resource-limiting level anticipated for these populations in the absence of phage, and free temperate phage are maintained by induction of lysogens at a density of P* = 2.5×10 ^4.^ In the absence of lysogens, sensitive cells S can be driven to extinction by clear phage P_C_ (Fig. 2C) (which would also be the case for lytic phage, P_V_ [28]). Both lysogens and sensitive cells can be driven to extinction by virulent phage, P_V_ (Fig 2D). This extinction can be attributed to a dearth of sensitive cells upon which the phage can replicate; once sensitive hosts are exhausted, the phage are diluted out at each transfer. However, when sensitive cells, N, are infected with both temperate phage P and clear mutants, P_C_, bacterial lysogens, N_L_ rapidly form and become the dominant population of bacteria (Fig. 1E). As a result of their adsorption to the lysogens and their inability to replicate on these lysogens, the density of clear mutants declines with the ascent of lysogens whilst the temperate phage are maintained.

**Fig 2.**
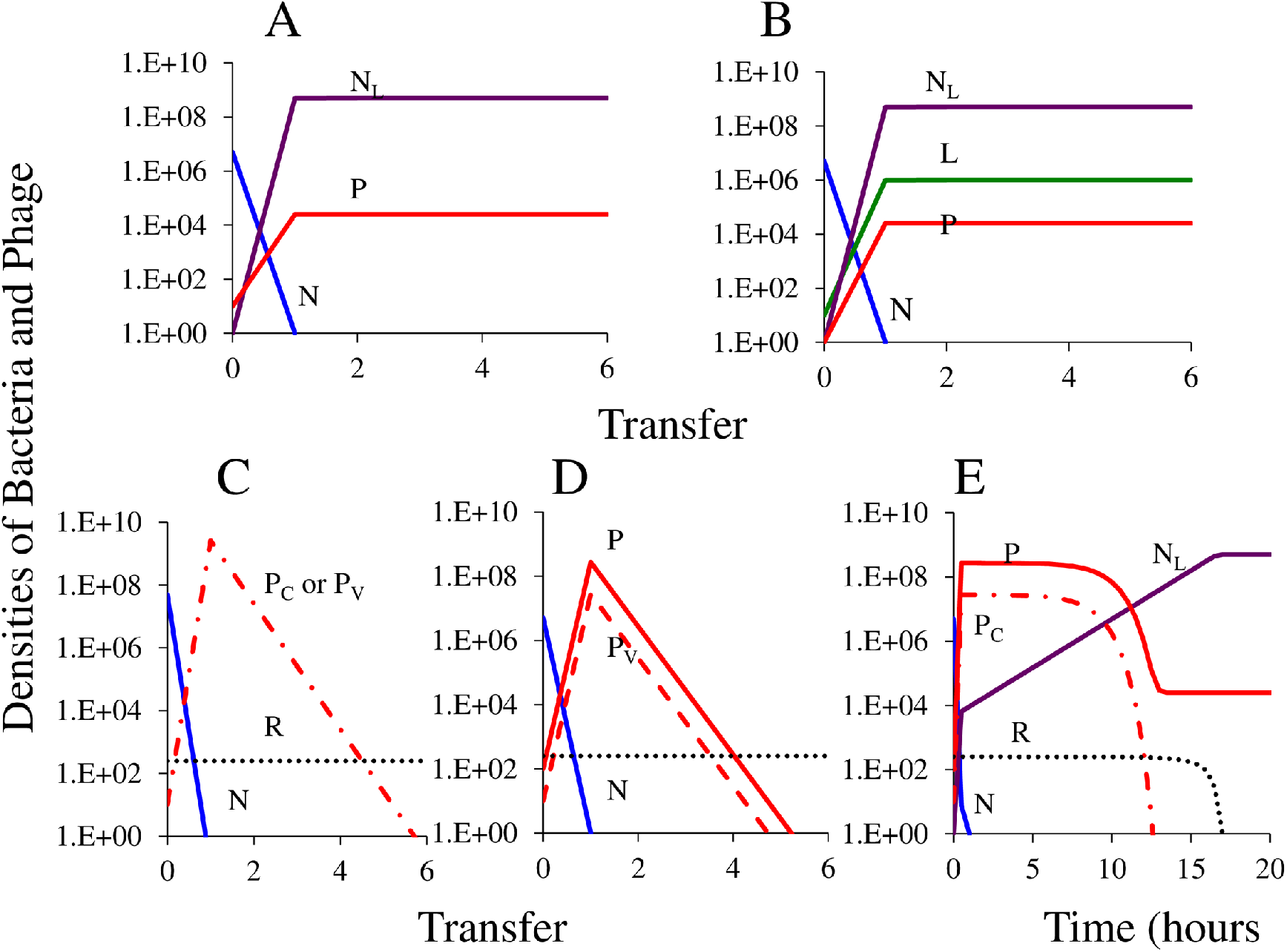
Population dynamics of lysogeny in the absence of resistance; densities of bacteria and phage at the start of the simulation and in (A)-(D) at the end of each transfer. Standard parameter values, vs = vr=0.7, k=1, e=5×10^−7^, δ=2×10^−7^, β=β_L_= 50, λ=0.001, R(0) =C=250, i=0.00001, μsr = μlsr = 0. (A) simulation initiated with 5×10^6^ sensitive cells, S, and 10 free temperate phage, P, (B) simulation initiated 5×10^6^ sensitive cells, S, and 10 lysogens, L. (C) simulation initiated with 5×10^6^ sensitive cells with 10 clear or virulent mutant phage, (D) simulation initiated with 10 virulent phage, P_V_, invading a population of lysogens N_L_ and free temperate phage at 1/100 of the equilibrium densities denoted in Fig 2A. (E) simulation initiated with 10 clear mutants, P_C_ and 10 free temperate phage, P, invading a population of sensitive non-lysogens in batch culture (time scale in hours).

#### 2-Simulated population dynamics with resistant mutants

When temperate phage invade a population of sensitive cells, most of the infections will be lytic, and resistant non-lysogens, N_R_ as well as immune lysogens, N_L_ will be favored by selection. If the rate of lysogeny is high enough, the density of lysogens will increase at a rate greater than that of these resistant cells, and lysogens will ascend to dominate the bacterial population. The resistant cells do not replace the lysogens and remain at relatively low densities (Fig 3A, 3B and 3C). A very different situation obtains when virulent mutants are present. Under these conditions, resistance engenders a considerable advantage. If the resistant cells are non-lysogens, N_R_, they become the dominant population of bacteria, and the temperate and virulent phage are both lost (Fig 3D). If the resistant cells are lysogens, N_RL_ the lytic phage are lost, but the population of free temperate phage continues to be maintained due to induction by the dominant population of resistant lysogens (Fig 3E).

**Fig 3.**
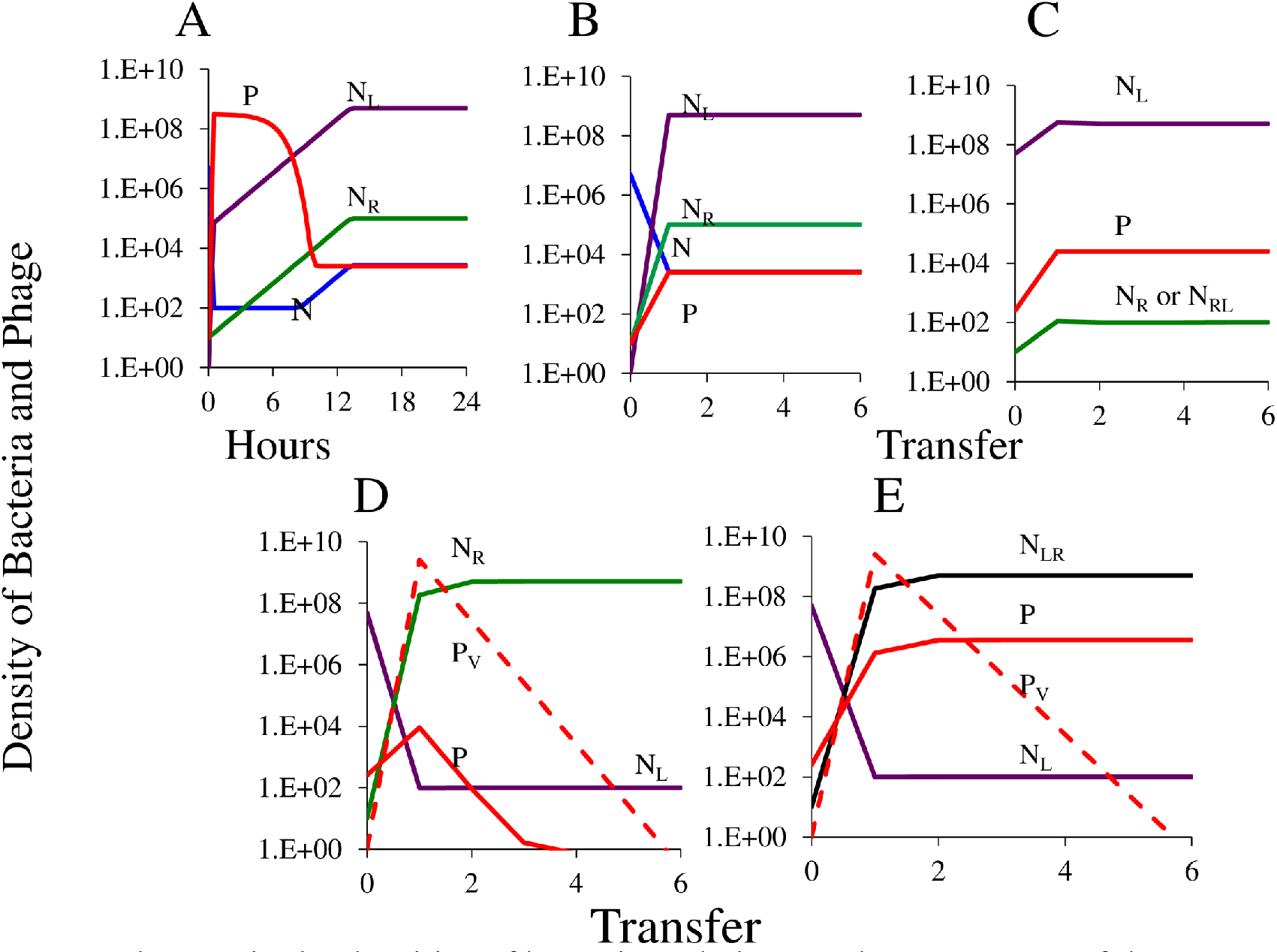
Changes in the densities of bacteria and phage. The parameters of these simulations are the same as those in Fig 2, save for a refuge where infection stops when the density of sensitive cells is less than 10^2^. (A, B) Simulation initiated with 5×10^6^ sensitive non-lysogens, S, 10 temperate phage, P, and 10 resistant non-lysogens, N_R_. (C) Simulation initiated with lysogens, N_L_, and free phage, P, at 1/100 of the at the equilibrium densities in Fig 2A and 10 resistant cells (N_R_ or N_RL_). (D) Simulation initiated with 1/100 of the equilibrium densities of P and N_L_ in Fig 2A plus 1 resistant non-lysogen, N_R_, and 1 lytic phage, PV. (E) Simulation initiated with 1/100 of the equilibrium densities of P and N_L_ in Fig 2A plus 1 resistant lysogen, N_RL_, and 1 lytic phage.

### II. Experimental Results

Arguably, surely in this case, mathematical models are more useful when they do not fit the data than when they do. That way they indicate that the biological assumptions upon which the models are based and/or the parameters used for the numerical analysis of their properties are in error. Ideally, these models will point to what has to be modified to better understand the biology under consideration. With this perspective in mind, we now explore the results of the experiments we have done addressing the different elements of the population dynamics of temperate and lytic phage considered in the previous section and thereby test the validity of the predictions generated from the model. These experiments are performed with *E. coli* MG1655 and a genetically marked temperate phage λ^KAN^, as well as clear and lytic mutants of this phage, λ^CL^, and λ^VIR^, in maltose-limited minimal medium. For details see the Experimental Methods.

#### 1-Establishment of temperate phage in populations of sensitive non-lysogens

In Fig 4 we present the results of experiments parallel to those modeled in Figs 2A and 2B. As anticipated by the model, when free temperate phage are introduced into a culture of sensitive non-lysogens, the culture rapidly becomes dominated by new lysogens (NL) which maintain a population of free temperate phage (P) (Figs 4A and 4B). While with our sampling procedure, we cannot rule out the possibility that there are a minority population of sensitive non-lysogens, it is clear that the dominant population of bacteria are newly generated lysogens. The densities of the free temperate phage in these experiments are similar to that anticipated by the model if the rate of induction is set at 1E-5 per cell per hour (Fig 4C). To independently explore the validity of this estimate of the induction rate, we established serial transfer cultures of resistant lysogens and estimated the densities of free phage at each transfer (Supplemental S1A Fig). As can be seen in the simulation presented in (S1B Fig), with an induction rate of 10^−5^ per cell per hour, the simulated density of free phage corresponds to that observed. A similar estimate of the induction rate for lambda lysogens was reported in [30].

**Fig 4.**
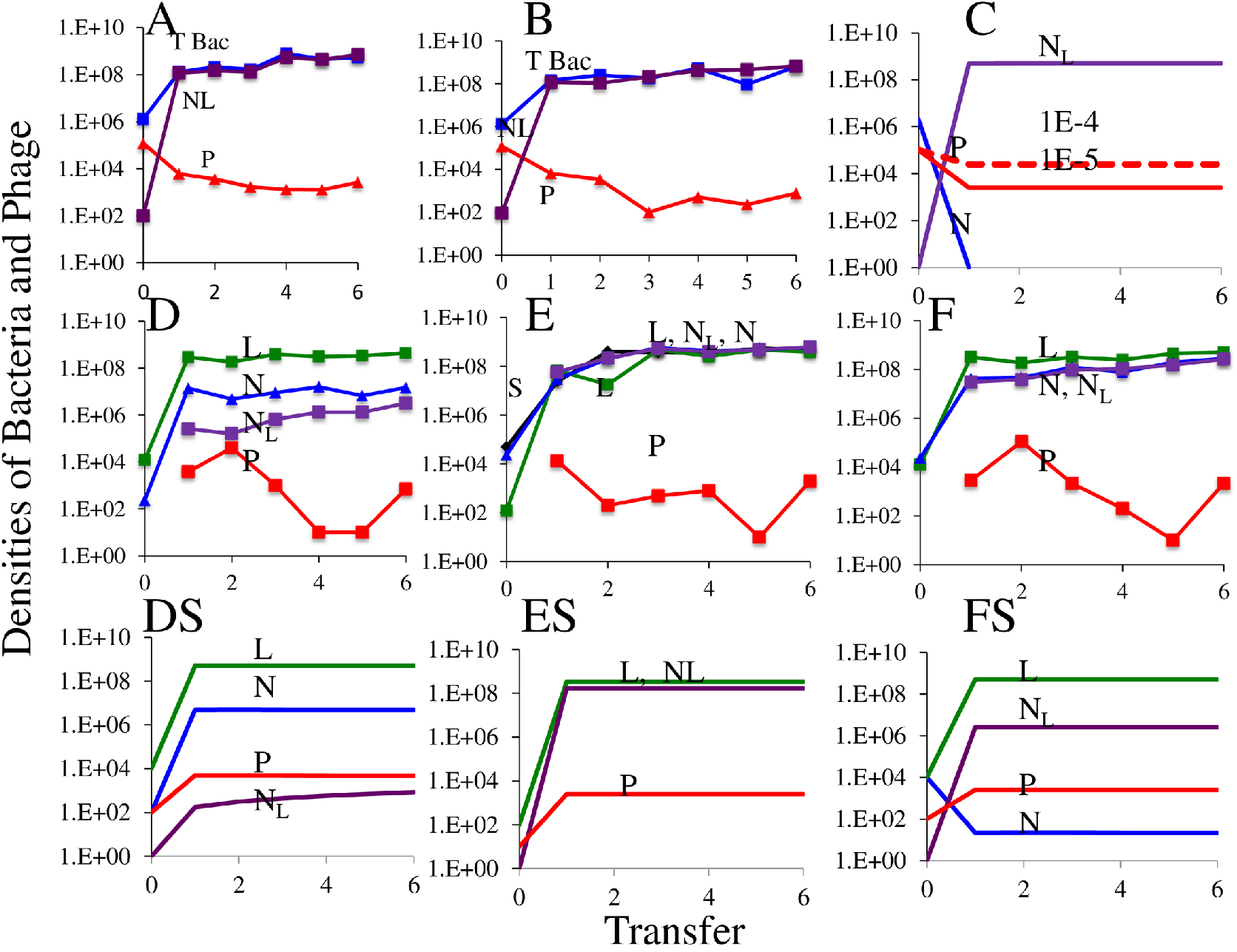
Establishment of lysogens in populations of sensitive non-lysogens. (A-C) Establishment of lysogens in populations of sensitive cells infected with free temperate phage. A and B show the results of two independent experiments, and C shows corresponding simulation results with the parameters used in Fig 2A, but with initial densities similar to that in the experiment and two different induction rates. (D-F) Establishment of new lysogens (Kan-R, Nal-R) and free temperate phage in populations with different initial densities of lysogens (Kan) and sensitive cells (Nal-R). DS, ES, and FS are corresponding simulation results with the initial densities similar to those in the experiments D, E, and F.

As anticipated from the model (Fig 2B), when populations of sensitive cells N are mixed with lysogens L, within short order free phage are produced and newly formed lysogens, N_L_, ascend to dominate the bacterial population. To better explore the dynamics of this situation, we considered three initial conditions: (i) Where the lysogens are a minority (Fig 4D), (ii) where the sensitive cells are the minority (Fig 4E) and (iii) where lysogens and sensitive cells are initially at the same density (Fig 4F). In Fig 4 DS, ES and FS, we present the results of the corresponding simulations. The relative densities of the original lysogens, L, newly generated lysogens, N_L_, and sensitive non-lysogens, N, in these simulations with different initial frequencies of L and N, are consistent with that observed. However, unlike the simulations, the densities of free temperate phage in the experimental cultures vary considerably in time for reasons that are currently unclear.

In accord with the simulation results presented in Fig 2B, if a population of temperate phage is mixed with sensitive non-lysogens, and if there is a minority of bacteria resistant to the phage, they will not ascend. To test this hypothesis, we mixed sensitive non-lysogens and free temperate phage in maltose limited minimal medium and added a minority population of bacteria resistant to both streptomycin and λ^VIR^, rpsL λ^VIR^. The results of this experiment are presented in Fig 5. The resistant non-lysogens became the dominant population, but newly formed lysogens also ascended and maintained densities about 20% less than the resistant (Figs 5A and 5B).

**Fig 5.**
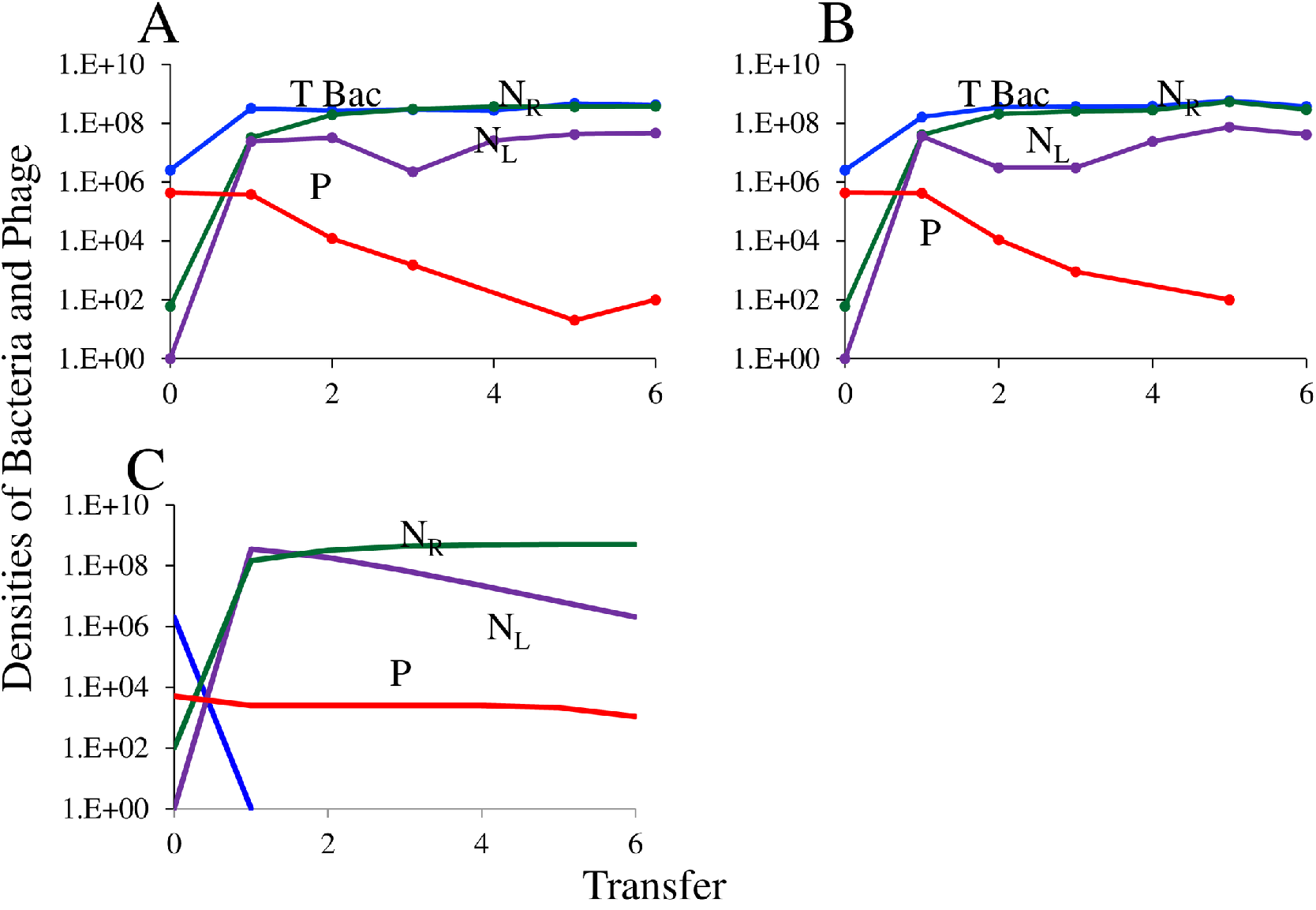
(A, B) Competition between sensitive N, and l resistant, N_R_ (Strep-R) in the presence of temperate phage P; data shown are two independent replicates. (C) Simulation initiated with sensitive N, temperate phage, P, and resistant non-lysogens, N_R_. Save for a higher growth rate for the resistant non-lysogens, vre =0.95 rather than 0.7, all other parameters are the same as in Fig 2A.

We can explain the results presented in Fig 5A and 5B by assuming that the Str-R l^VIR^ resistant are more fit than the sensitive non-lysogens, see the simulation results in Fig 5C. If the maximum growth rate is a reflection of relative fitness, this is indeed the case (S2 Fig). As the Str-r, *rpsL* mutation borne by l^VIR^ – resistant cells (an arginine (R) replacing the wild type lysine (K) at the 43^rd^ codon) has no measurable fitness cost or benefit [31], the resistance marker does not account for the observed fitness difference. At this juncture we don’t know why these l-resistant non-lysogens are more fit than the wild type sensitive cells.

#### 2-Dynamics of clear mutant, temperate phage, and sensitive bacteria

In Fig 6 (A, B, C) we follow the changes in densities of bacteria and phage in a serial transfer population initiated with sensitive cells and a clear mutant phage. Contrary to what is anticipated from the corresponding theoretical result, Fig 2C, the sensitive bacterial population is not eliminated following the ascent of λ^CL^, but rather is maintained. We postulate that this deviation between theory and experiment can be attributed to the generation of temperate λ by the clear mutants, and subsequently by the conversion of the sensitive cells into lysogens [32] and selection for and ascent of envelope resistant non-lysogens. There are three lines of evidence in support of this hypothesis. 1-There were turbid plaques on the plates estimating the densities of free phage, consistent with plaque morphology for temperate λ. 2-When exposed to UV, the purified colonies from sixth transfer of the experiment Fig 6A produce free phage, consistent with the induction of lysogens. Interestingly, while the original temperate phage λ^KAN^ conferred resistance to kanamycin in its host, the reverted clear phage derived from this temperate strain had lost or inactivated this drug-resistance cassette and no longer conferred resistance to the drug. 3-The purified colonies from the sixth transfer of the experiment Fig 6 (B, C) are resistant to λ^−VIR^.

**Fig 6.**
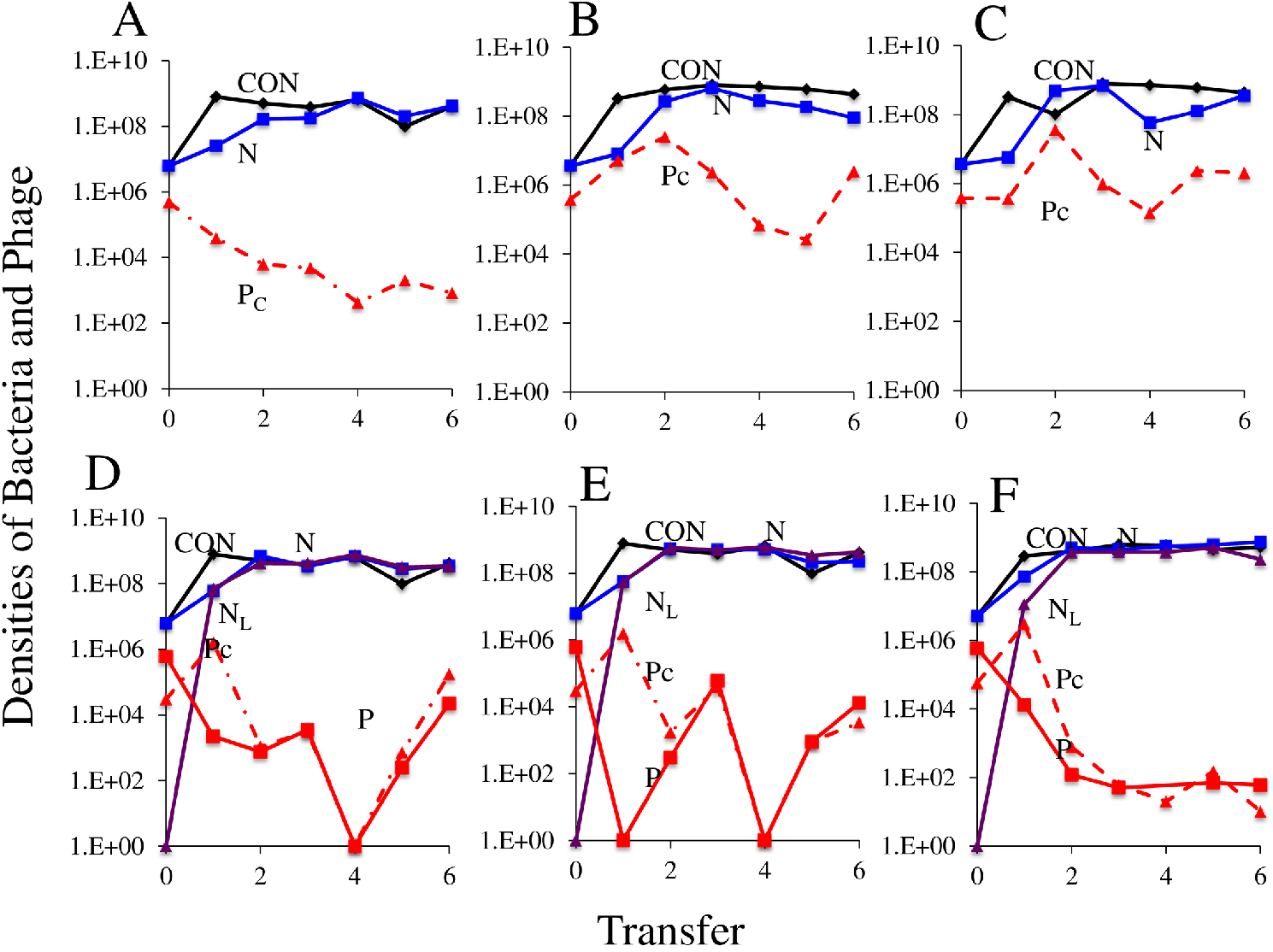
Population dynamics of sensitive bacteria with λ clear mutant phage. Data shown are the densities of bacteria and phage over six serial transfers. A, B, C-Cultures initiated with clear mutant phage (P^C^) and sensitive bacteria, N. (D, E, F)-Cultures initiated with temperate (P) and clear (P^C^) phage and sensitive bacteria, N.

The clear mutant in Fig 6A reverted back to the temperate mode by excising out single base insertion mutation at position 673 in cI repressor. The sequences of *cI* gene of wild type λ, clear mutant, and their revertants have been submitted to GenBank with accession number MN510929, MN510930 and MN510931, respectively. While we would certainly expect clear mutants to revert to temperate but why would the temperate phage take over? A likely answer is that both the temperate phage and the clear mutants are lost when they adsorb to the lysogens. This can be seen in Fig 6 (D, E, F) where temperate phage P and clear P^C^ mutants are introduced together in a culture of sensitive cells.

When mixed with λ^+^ and sensitive cells, lysogens evolve to dominate the culture, but clear and temperate phage continue to coexist (Fig 6 D, E and F)., contrary to the predictions of the model (S3 Fig). This occurs despite the fact that the clear phage are unable to replicate on lysogens (S4 Fig). On the other hand, A possible explanation for this was presented by Armin Kiaser in 1957 [33] by a mechanism he called “double lysogeny”. When sensitive cells are simultaneously infected by temperate l and clear mutants, both became integrated into the chromosome. Upon induction, these double lysogens produced clear as well as temperate phage.

#### Dynamics of virulent mutant, temperate phage, and sensitive bacteria

In Fig 7 we present population dynamics results of virulent mutant phage and temperate phage with sensitive bacteria. The resistant non-lysogen evolved to dominance by first transfer, while the newly formed lysogens were present as a sub-population. Both lytic and temperate phage were maintained in these cultures. At one level, the results of the experiments presented in Fig 7A and 7B are consistent with that anticipated from the corresponding simulation in Fig 3D, mutants with envelope resistance, N_R_, emerged and ascended to dominate the bacterial population. Inconsistent with the simulation is the continued maintenance of both temperate and lytic phage, P and P_V_ and a minority population of lysogens, N_L_. Included among the cells in the N_L_ population were those with envelope resistance for the lytic phage, resistant lysogens, N_LR_. By selecting for kanamycin resistance, the relative frequency of lysogens increased (Fig 7C). However, contrary to what is anticipated from this model in Fig 3D, in all three cultures both the temperate and lytic phage are maintained, rather than lost. We postulate that these phage are maintained by the leaky resistance mechanism we demonstrated for λ^VIR^ and *E. coli* K12 [28], primarily transitions from N_R_→S and N_LR_→N_L_.

**Fig 7.**
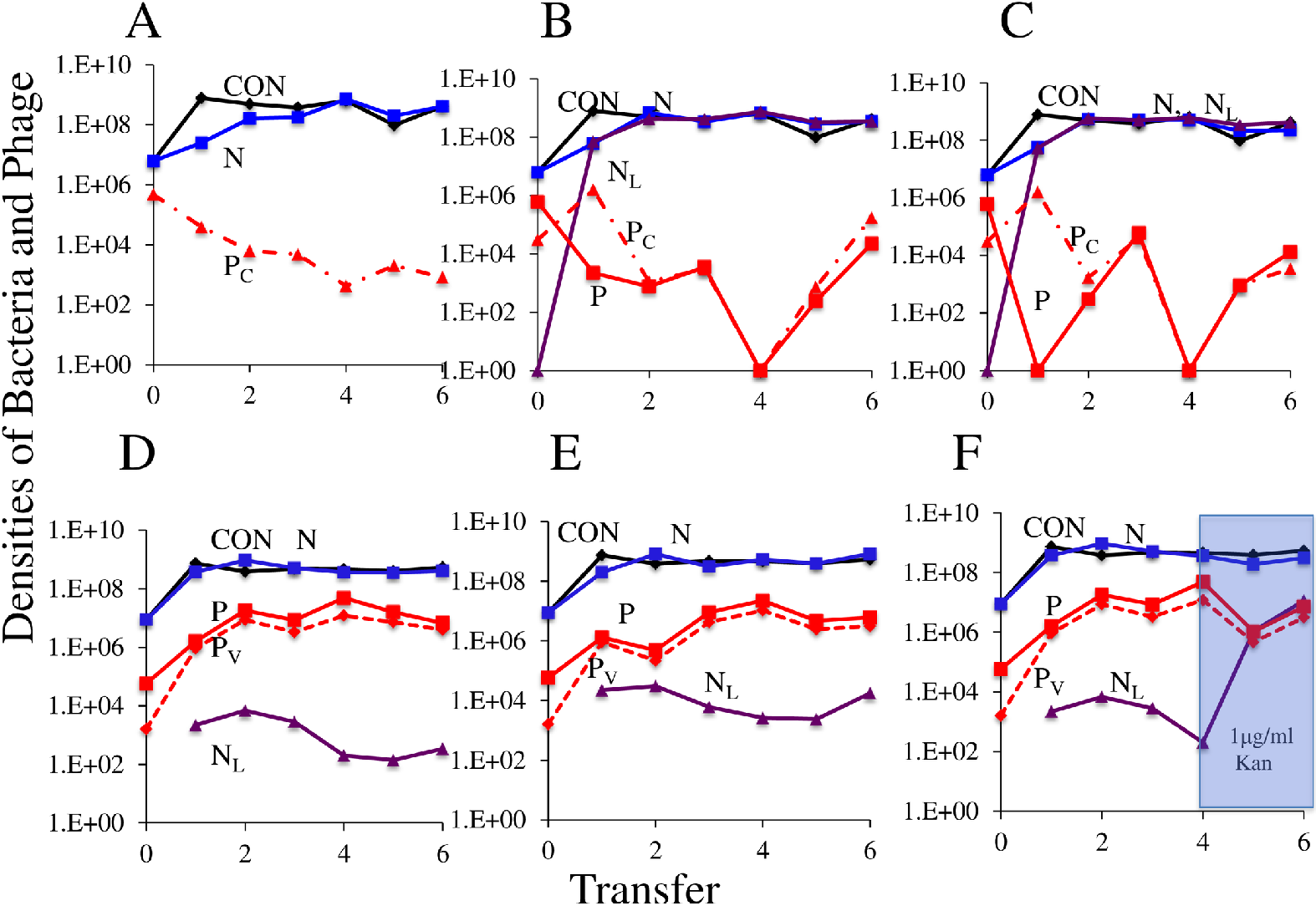
Population dynamics of virulent mutant, temperate phage and sensitive bacteria. (A, B, C) Cultures initiated with temperate P and virulent phage, P^V^, and sensitive cells, N. The serial transfer culture depicted in Fig C was produced by adding 1μg/ml of kanamycin to the medium at the 4th transfer of the experiment depicted in Fig B. (D, E) Cultures initiated with temperate, P, and virulent phage, P_V_ and sensitive lysogens, L, and minority population of resistant lysogens N_RL._ (F) Simulation results of temperate, virulent phage and sensitive lysogen, L, with resistant lysogens NLR, vl=1.0^5^ and transitions between states, μsr=μrs= 5×10^−5^

In Fig 7(D, E) we consider what obtains in the reconstruction experiment when resistant lysogens, N_RL_, are present as a minority along with temperate and lytic phage and majority sensitive lysogens. In all replicas, the resistant lysogens ascend and become the dominant population of bacteria. In three of the five replicas, both temperate and lytic phage were still present at the 6^th^ transfer. Maintenance of the virulent phage would be anticipated if the rate of transition from resistant non-lysogens is to sensitive cells, NR → S, μrs, is high enough to maintain the phage on a minority population of sensitive lysogens (Fig 7F). However, this loss of the temperate phage is inconsistent with that anticipated from the model (S5 Fig).

## Discussion

In an article on the population dynamics and evolution of lysogeny published before the majority of authors and readers of this report were diploid, Stewart and Levin [5] called for experimental studies to test the hypotheses generated from their theoretical analysis. This investigation tests those hypotheses with *E. coli* K12 and the phage l. Although the model we use here is based on that by Stewart and Levin [5], it differs from this model and other more recent models addressing the population dynamics of temperate phage [4, 34–38] in four ways; (i) it considers clear mutants, which can infect sensitive cells but not form or replicate on lysogens, (ii) it allows for resistant non-lysogens as well as resistant lysogens which maintain the prophage but are refractory to both lytic and temperate phage, and (iii) as in [28] it allows for transitions between the susceptible and resistant states, and (iv) it considers populations maintained in serial transfer rather than continuous culture habitat. Moreover, and most importantly, to our knowledge this is the first experimental study of the population and evolutionary dynamics of temperate phage to consider virulent and clear mutants of these viruses and bacteria resistant to these temperate phages.

To a gratifying (surprising?) extent, the majority of the results of our experiments are, at a qualitative (semi-quantitative?) level consistent with the prediction generated from our analysis of the properties of our extended version of the Stewart - Levin Disco-era model. As anticipated by the model, when free temperate λ^+^ are introduced into populations of sensitive non-lysogens, within a single transfer newly formed lysogens become the dominant population of bacteria [20, 35]. Free temperate λ^+^ are maintained at densities in a range anticipated from the burst size and the rate of induction of lysogens. As anticipated from this model, when λ^+^ lysogens are introduced into a population of sensitive cells, free phage λ^+^ are generated by induction and new lysogens are produced [35]. The relative densities of new lysogens, original lysogens and sensitive cells in these cultures, depends on initial densities of the original lysogens and sensitive cells in a way that is consistent with that predicted by the model and previous studies[39] With a single mutation in the *malT* or *lamB* genes, sensitive *E. coli* can become resistant to λ^+^ as well as its clear and virulent mutants [28, 40, 41]. Envelope resistance via these mutations confers a fitness cost in the minimal maltose medium [42]. When temperate phage invade a population of sensitive cells, because most of the infections are lytic, they would select for resistant mutants. Our model suggests that while this is the case, because lysogens are immune to free phage, under broad conditions rare populations of resistant cells will not ascend and will be washed out from the culture if they are less fit (Fig 5 D). They can only ascend if they are more fit than the lysogens, (Fig 5 D), which is very unlikely.

During the early phase of λ temperate phage infection, the density of susceptible cells is large (Fig 4A, 4B) and the free phage have opportunity for horizontal transmission by lytic infection [4]. λ phage lytic infection selects for envelope resistant bacteria which are less fit in the initial serial transfers of cultures initiated with sensitive bacteria and λ lytic phage, as can be seen in Fig 3 C and D in [28]. If the rate of lysogeny is high enough, the density of lysogens will increase at a rate greater than that of these resistant cells, and ascend to dominate the bacterial population (Fig 4A, 4B). Our reconstruction experiment with minority populations of l-resistant bacteria, high densities of sensitive cells and λ phage is in agreement with this hypothesis. However, the envelope resistant mutant bacteria used in the reconstruction experiment are more fit than the wild type. If the resistant bacteria are more fit or equally fit, and rate of lysogeny is low because of more lytic infections, the resistant bacteria will ascend to dominate the bacterial population. On the other hand, if lytic phage are present, our models predict that resistant mutants ascend and become the dominant population of bacteria. Unless other processes are in operation, the ascent of these resistant mutants will lead to the extinction of the sensitive lysogens and sensitive cells and thereby the loss of the phage. However, if those “other processes” include some mechanism for the generation of sensitive cells, like leaky resistance [28], the sensitive cells and/or lysogens can be maintained along with lytic and free temperate phage. At a qualitative level, the results of our experiments are consistent with this prediction. When serial transfer cultures are initiated with mixtures of sensitive *E. coli*, virulent and temperate phage, λ^VIR^ and λ^+^ resistant mutants emerge and ascend to dominate the bacterial population, but both temperate and lytic phage continue to persist along with a minority population of newly formed lysogens. The frequency of these newly formed lysogens is markedly increased if, the addition of low concentrations of kanamycin, which selects for the carriage the λ^KAN^ prophage [19, 43, 44] A recent study showed that temperate phage of *P. aeruginosa* in the lungs of CF patients carries antibiotic resistance genes and relates to the progression of the disease by providing a selective advantage to their host bacteria [45].

Most importantly, the results of these experiments with combinations of lytic and temperate λ indicate that a substantial fraction of the resistant bacteria evolving in these populations are lysogens. As a consequence, even if sensitive populations of lysogens are eliminated, the temperate phage will be maintained as a prophage and, by induction as free phage. To our knowledge, this is the first time the importance of resistant lysogens in the population and evolutionary dynamics of lysogeny has been reported. This observation also leads to a testable and broadly important prediction. If lytic phage are generated by mutation in natural populations of temperate phage, or those communities include lytic phage with the same receptors as the temperate, lysogenic bacteria isolated from natural populations will be resistant to the phage generated by induction. If indeed this is the case, phage induced from naturally occurring lysogens will not be able to adsorb to these lysogens.

### Caveats and excuses

As pleased as we may be about the relatively good fit between theory and experiments, we are very much aware of the limitations of our models and experiments. Most importantly, they are restricted to mass action models and the experimental analog, well agitated liquid cultures of bacteria. To be sure, there may well habitats the real world where bacteria exist as planktonic cells frolicking about in liquid all with equal access to resources and infecting phage. We expect that more commonly natural populations of bacteria and phage exist in physically structured habitats on surfaces, in semisolids or in biofilms along with other bacterial species and phages [46]. Moreover, it seems unlikely, that in natural populations of bacteria are as well or as reliable fed as they are in our models and experiment, but rather live in feast and famine conditions [47]. At this juncture, how these realities of bacterial life in the real world affect the population and evolutionary dynamics of bacteria and their temperate and lytic phage, are not at all clear, but can be addressed experimentally if not with simple mathematical models.

The molecular geneticists Jacque Monod is reported to have quipped, ‘*what is true for E. coli is true for Elephants*’. As appealing as this coli-centric form of inductive inference and it’s extension to the phage Lambda are, we are very much aware that the results reported here may not reflect that which obtains in with other bacteria and their temperate phage. It will be of some interest to address the “why be temperate question” with other bacteria and their temperate phage.

## Material and Methods

### Strains and growth media

All *E. coli* strains used in our experiments were derivatives of the parent strain K12 MG1655. The parent strain *E.coli* MG1655 was obtained from Ole Skovgaard at Roskilde University in Denmark. The λ^VIR^ strain used in these experiments was obtained from Sylvain Moineau at Laval University Quebec, Canada. The construction of temperate phage λ^KAN^ is described in [30]. λ^Clear^ was generated from λ temperate by spontaneous mutation.

Bacterial cultures were grown at 37°C either in M9 media [M9 salts (248510, Difco) supplemented with 0.4% maltose (075K0123, Sigma), 1 mM MgSO_4_ (Sigma Aldrich), 0.1 mM CaCl_2_ (JT Baker), and 0.2% Thiamine B1 (Sigma Aldrich)] or LB broth [MgSO4 2.5g/L, tryptone (Fisher Bioreagent 10g/L, yeast extract (Bacto) 5g/L, sodium chloride (Fisher Chemical) 10g/L]. λ temperate phage lysates were prepared from single plaques at 37°C in M9M medium alongside wild-type *E.coli* MG1655 by plate lysis. Specifically, individual phage plaques were picked with a sterile bamboo stick, resuspended in 3 ml of phage soft agar together with 100 μl of overnight bacterial culture and plated on top of phage plates. The plates were then incubated at 37 °C overnight. The soft agar was scraped with a sterile iron scoop, resuspended in 10 ml of M9 with a few drops of chloroform to kill the residual bacteria. The lysates were then centrifuged to remove the agar, sterilized by filtration (0.2 μm) and stored at 4 °C.

### Sampling bacterial and phage densities

Bacteria and phage densities were estimated by serial dilutions in 0.85% NaCl solution followed by plating. The total density of bacteria was estimated on LB (1.6%) agar plates. To estimate the densities of λ^KAN^ lysogens, cultures were plated on LB (1.6%) agar with 25 μg/mL kanamycin (AppliChem Lot# 1P0000874). To estimate the densities of free phage, chloroform was added to suspensions before serial dilutions. These suspensions were mixed with 0.1 mL of overnight LB grown cultures of wild-type MG1655 (about 5×10^8^ cells per mL) in 3 mL of LB soft (0.65%) agar and poured onto semi-hard (1%) LB agar plates.

### Bacteriophage and bacteria parameter determination

The parameters critical for the interaction of λ phages and *E. coli* MG1655 used in this study were estimated in independent experiments M9M. The maximum growth rate of different clones of *E.coli* MG1655 was measured by Bioscreen as described in [48]. Phage burst sizes (β) were estimated with one-step growth experiments similar to [49]. Adsorption of λ to *E. coli* was estimated as described in [49]. The procedure for estimating the probability of lysogeny and the rate of spontaneous lysogen induction are presented in [30].

### Short term dynamics experiments

Overnight cultures of *E. coli* MG1655 strain in M9M medium were grown. The cultures were diluted in a 1:1000 ratio on 20 mL of medium in 100 mL flasks and were incubated for 4 hours to achieve a density of 1×10^6^ CFU/mL and λ^KAN^ phage was added to an initial density of 1×10^5^ PFU/mL. Short term dynamics were conducted culturing the mixture at 37 °C in shaking conditions. Total bacteria count was conducted by sampling 100 μL at regular intervals and plating dilutions on LB agar plates. λ^KAN^ lysogens were obtained by sampling 100 μL and plating dilutions on LB agar with 25 μg/mL kanamycin plates. Phage count were obtained by sampling 1 mL at regular intervals and adding.

### Serial Transfer

All serial transfer experiments were carried out in 10 ml M9M cultures grown at 37°C with vigorous shaking. The cultures were initiated by 1:100 dilution from 10-mL overnight cultures grown from single colonies. Phage were added to these cultures to reach the initial density of approximately 10^5^ PFU/mL. At the end of each transfer, 100 μL of each culture was transferred into flasks with fresh medium (1:100 dilution). Simultaneously, 100 μL samples were taken for estimating the densities of colony forming units (CFU) and plaque forming units (PFU), by serial dilution and plating on solid agar, with selection as needed as described above.

### Identifying mutation for λ_CL_ and λ_CL_ revertant

The cl gene of the wild type λ temperate, clear mutant and clear revertant mutant phage was amplified from the individual plaques using the fw-cI (TTGCTGCGGTAAGTCGCATA) and rv-cI primers (GCGCTTTGATATACGCCGAG) in a standard PCR reaction with Taq DNA polymerase (Thermo Scientific Phusion High-Fidelity DNA Polymerase). The PCR products were then purified using the GenElute Gel Extraction kit (Sigma Lot# SLBL6694V) and sequenced. Mutation were identified by comparing the sequences of the mutants to the corresponding sequence of wild type λ temperate

## Acknowledgements

We are grateful to Maros Pleska and Calin Guet for providing us with the phage used in this study and for insightful comments Maros Pleska provided during the early stage of this investigation and Joshua Weitz. This research was supported by a grant from the National Institutes of Health, GM091875 (BRL), Emory University Start-up funds (NV) and Beca CONICYT PFCHA DOCTORADO/2016 21161133 funds to Rodrigo Garcia to support his visit to Emory University.

This report is dedicated to the memory of Allan M. Campbell and Frank M. Stewart and the inspiration they provided to BRL and by induction the other authors.

---

**S1 Fig.**
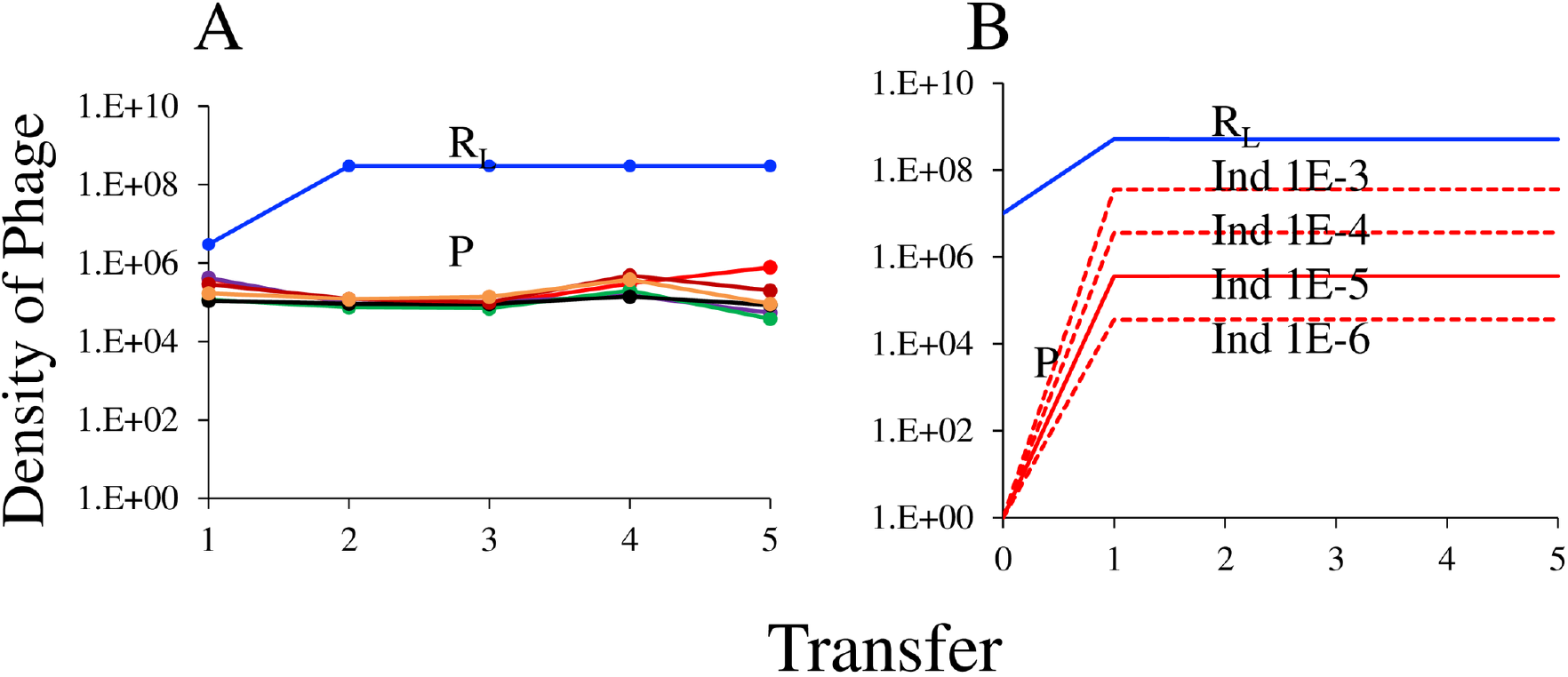
(A). The densities of free phage in the serial transfer culture of six independently selected MG λ VIR resistant lysogens, N_RL_, in M9-maltose. (B) Corresponding simulation results with different induction rates for resistant lysogens.

**S2 Fig.**
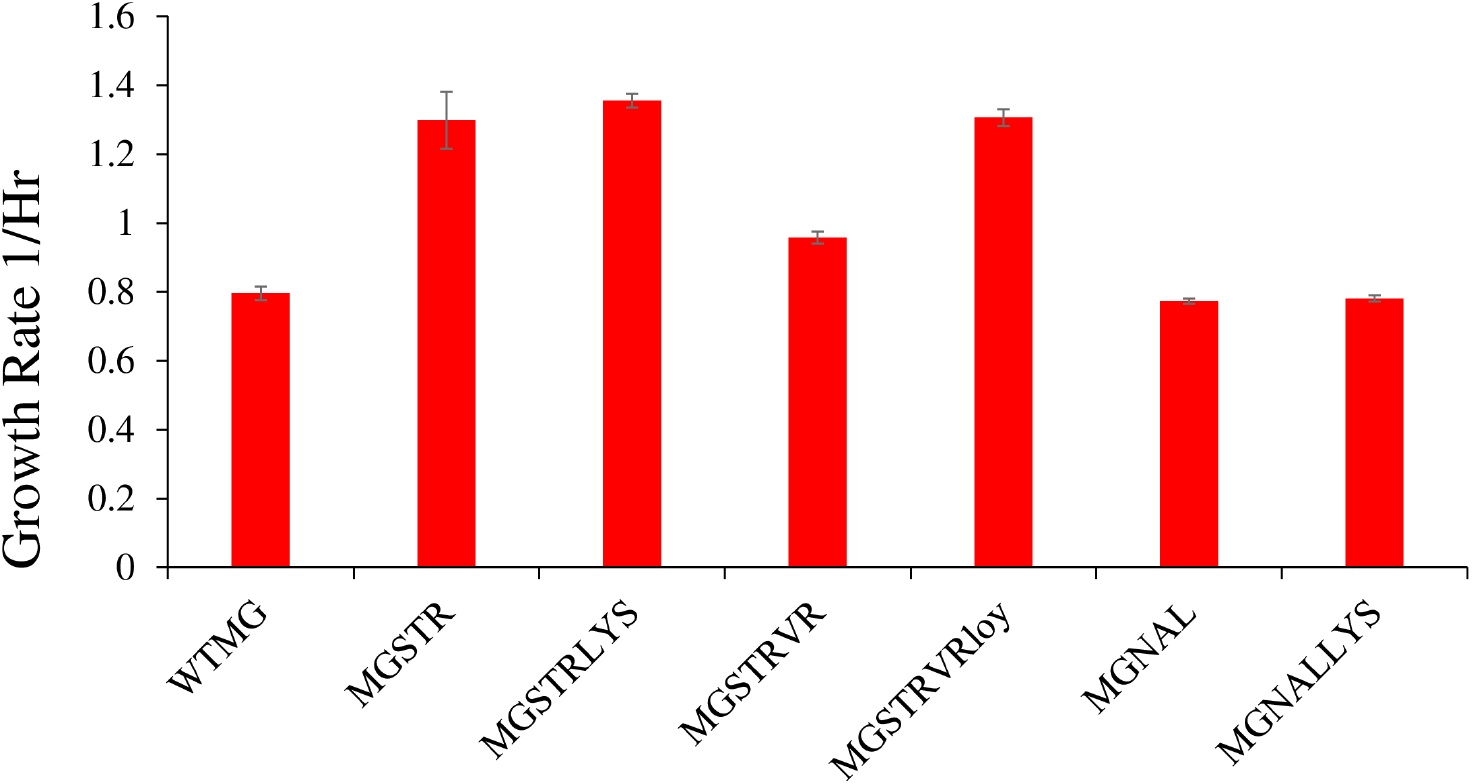
Estimated growth rate. Data were obtained from OD_600_ growth curves in M9-maltose at 37°C. WT MG: WT *E.coli* MG1655, MGSTR: *E.coli* MG1655 resistant to Streptomycin, MGSTRLYS: *E.coli* MG1655 resistant to Streptomycin and lysogen with lambda temperate (kan) phage, MGSTRVR: *E.coli* MG1655 resistant to Streptomycin and resistant to lambda vir phage, MGSTRVRLoy: MGSTRLYS resistant to lambda vir phage, MGNAL: *E.coli* MG1655 resistant to Nalidixic acid, MGNALLYS: *E.coli* MG1655 resistant to Nalidixic acid and lysogen with lambda temperate (kan) phage.

**S3 Fig.**
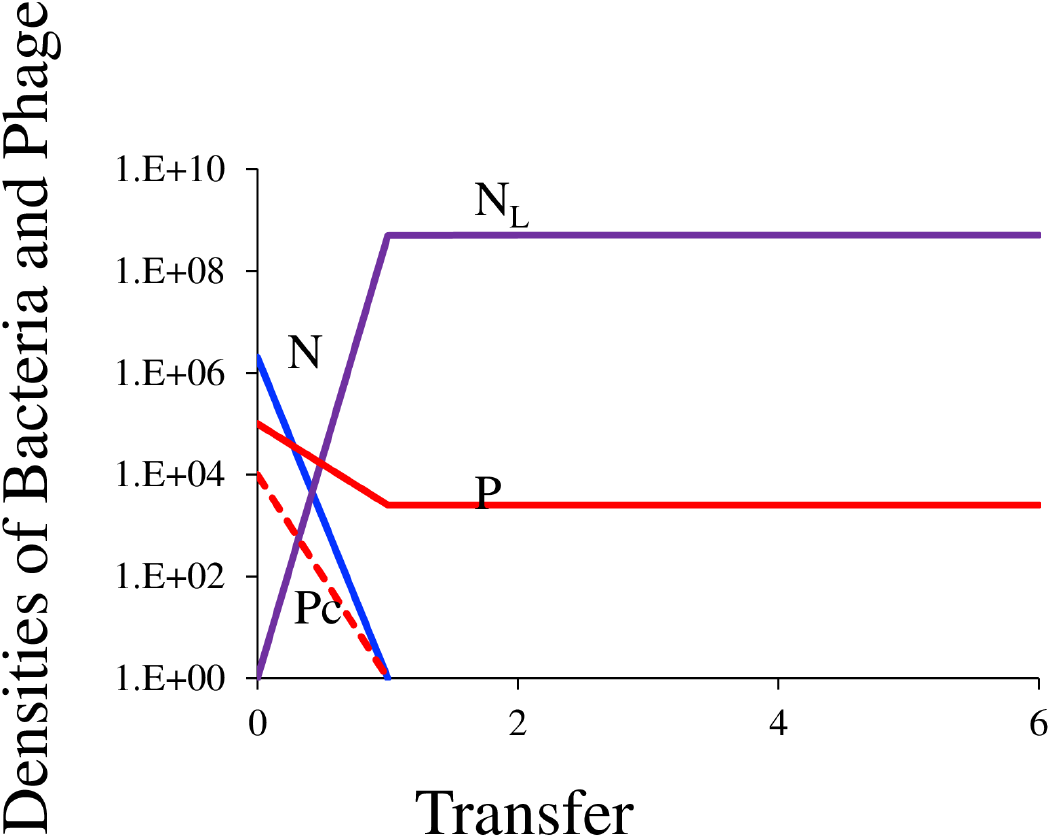
Simulation initiated with sensitive, N, temperate phage, P, and clear mutant, Pc.

**S4 Fig.**
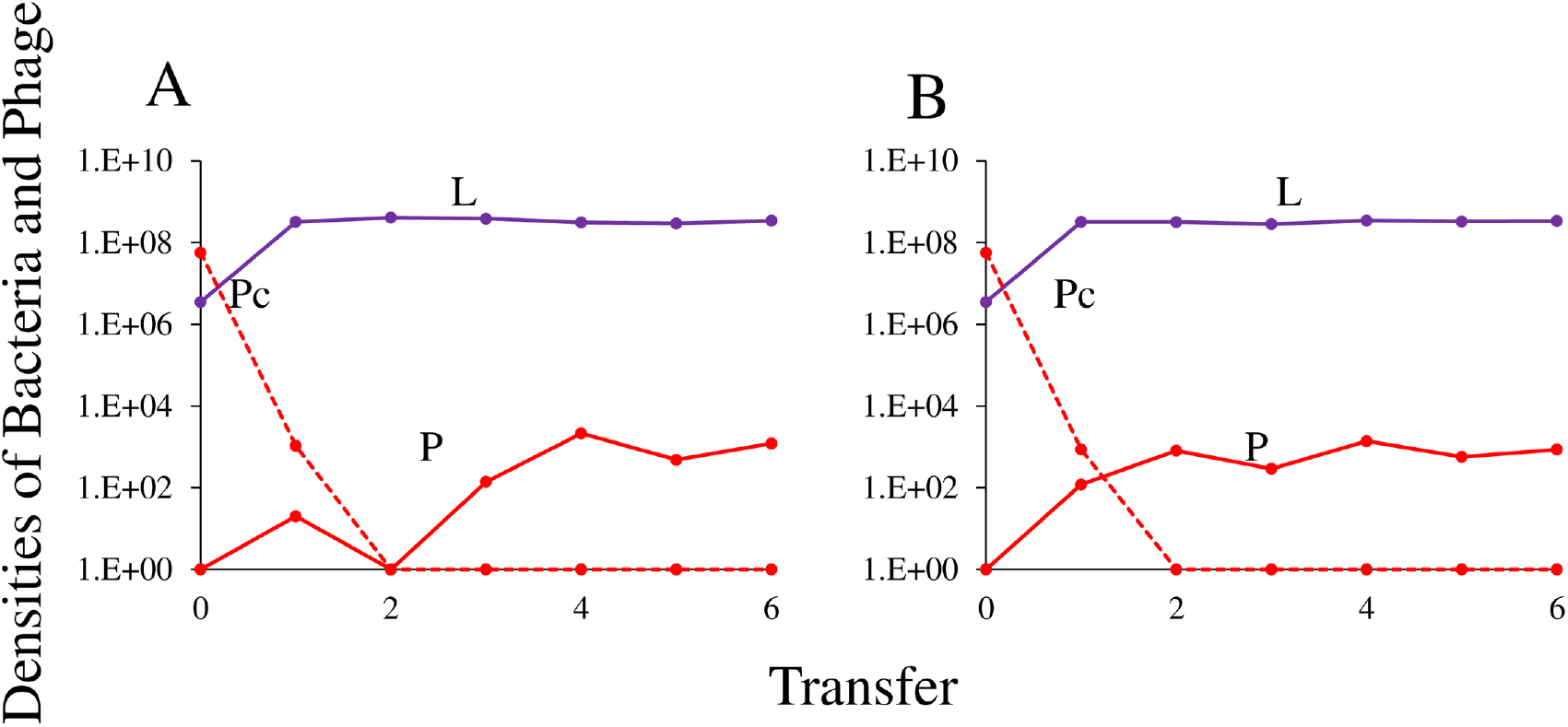
Population dynamics of Lysogen with λ clear mutant phage in two independent replica experiments (A, B).

**S5 Fig.**
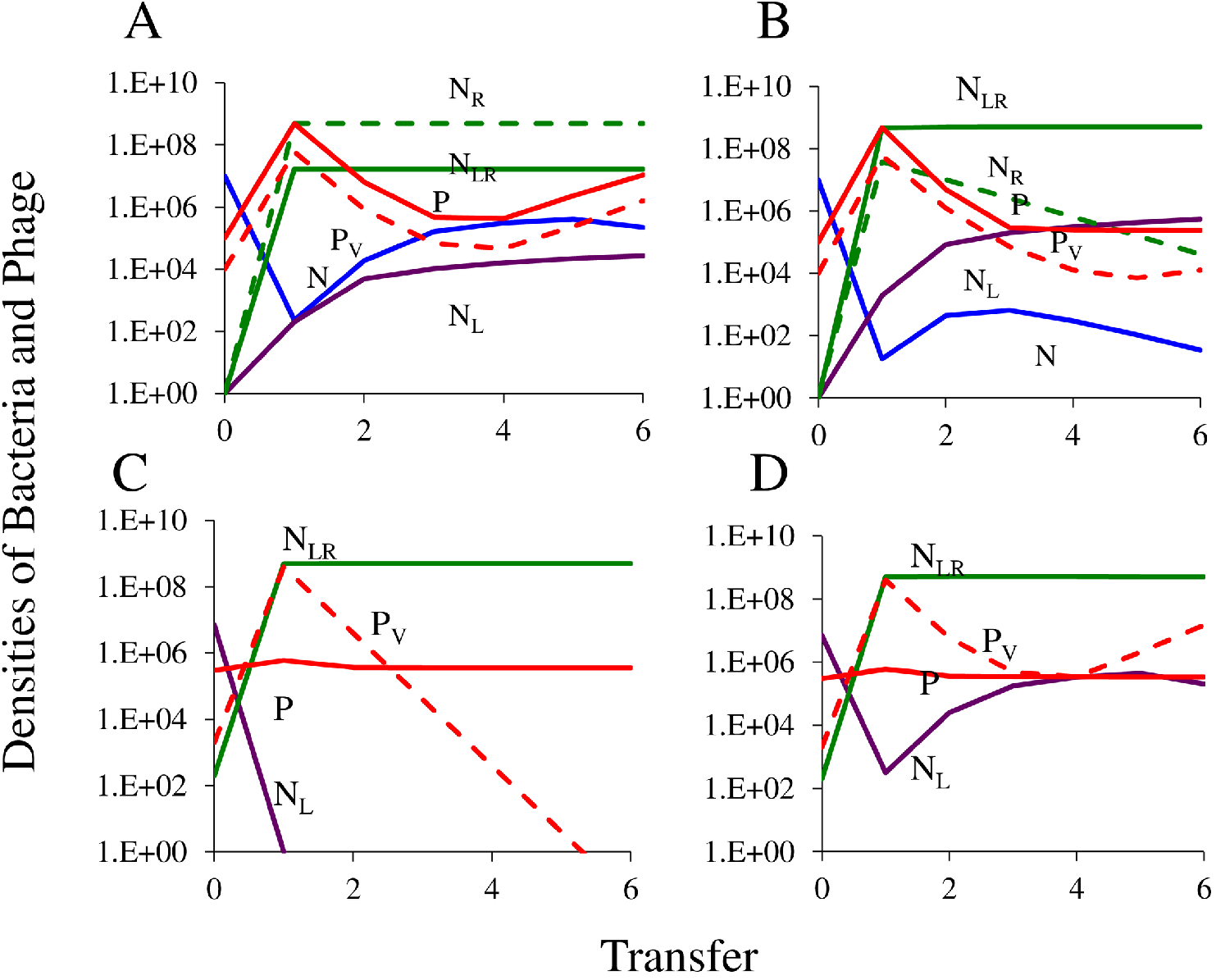
Simulated population dynamics, changes in the densities of bacteria and phage. Standard parameters: v = vl =vr = 0.7, δ=2×10^−7^, β=50, β_L_=50, λ=0.01, i=10^−5^, C=250, e=5×10^−7^, k=1.0, d=0.01. (A) Temperate and virulent phage, P and P_V_ and sensitive non-lysogens, N, with single resistant non-lysogens N_R_ and lysogens NLR, equal fitness for all bacterial populations, with transitions between states μsr=μrs= 5×10^−5^. (B) Temperate and virulent phage, P and P_V_ and sensitive non-lysogens, S, with single resistant non-lysogens and lysogens, with fitness advantage for lysogen, NL and NLR, vl=1.0^5^ and transitions between states, μsr=μrs= 5×10^−5^ (C) Temperate and lytic phage introduced into a population of sensitive, N_L_ and resistant lysogens, N_RL_, no transitions between states, μsr=μrs=0 (D) Temperate and lytic phage introduced into a population of sensitive, N_L_ and resistant lysogens, N_RL_, with transitions between resistant and sensitive states, μrs=μsr=5×10^−5^.

